# Dissection of prostate tumour, stroma and immune transcriptional components reveals a key contribution of the microenvironment for disease progression

**DOI:** 10.1101/2020.03.16.993162

**Authors:** Stefano Mangiola, Patrick McCoy, Martin Modrak, Fernando Souza-Fonseca-Guimaraes, Daniel Blashki, Ryan Stuchbery, Simon P. Keam, Michael Kerger, Ken Chow, Chayanica Nasa, Melanie Le Page, Natalie Lister, Simon Monard, Justin Peters, Phil Dundee, Anthony J. Costello, Paul J. Neeson, Scott G. Williams, Bhupinder Pal, Nicholas D. Huntington, Niall M. Corcoran, Anthony T. Papenfuss, Christopher M. Hovens

## Abstract

Prostate cancer is caused by genomic aberrations in normal epithelial cells, however clinical translation of findings from analyses of cancer cells alone has been very limited. A deeper understanding of the tumour microenvironment is needed to identify the key drivers of disease progression and reveal novel therapeutic opportunities. In this study, the experimental enrichment of selected cell-types and the development of a Bayesian inference model for continuous differential transcript abundance permitted us to define the transcriptional landscape of the prostate cancer microenvironment along the disease progression axis. An important role of monocytes and macrophages in prostate cancer progression and disease recurrence was uncovered, supported by both transcriptional landscape findings and by differential tissue composition analyses. These findings were corroborated and validated by spatial analyses at the single-cell level using multiplex immunohistochemistry. This study advances our knowledge concerning the role of monocyte-derived recruitment in primary prostate cancer, and supports their key role in disease progression, patient survival and prostate microenvironment immune modulation.

## Background

Prostate cancer is the second most commonly diagnosed cancer in men globally^1^. Although most cancers follow an indolent clinical course, an unpredictable 10-15% of tumours progress to metastases and death. The inability to discern progressive disease at an early stage leads to substantial overtreatment of localised disease, leading to a significant clinical cost to the patient and economic cost to the healthcare system. Selecting patients for treatment is usually reliant on a small number of well-established clinical and pathological factors, such as tumour grade, prostate serum antigen (PSA) level and clinical stage, which have been consistently associated with disease recurrence^2^, the development of metastases^3^ and prostate cancer-specific death^4^. Although comprehensive molecular analyses have linked clinical outcomes with rates of genomic alterations^5,6^, such as somatic changes in copy number, nucleotide sequence and methylation, it has yet to be demonstrated that such measures are able to consistently outperform standard clinico-pathological risk scoring across a broad range of grades and stages. Despite many years of tumour evolution characterization, it still remains unclear what mechanisms drive prostate cancer progression in most patients^7^. It is believed that reciprocal interactions between malignant epithelium and surrounding non-cancerous cells within the tumour microenvironment are responsible for driving disease progression^8,9^.

Selected targets in the prostate tumour microenvironment have been extensively studied through *in vitro* and *in vivo* experiments, such as migration assays^10^ and xenograft mouse models^11^ respectively. More recently, several studies that integrated fluorescence-activated cell sorting (FACS) or laser microdissection with RNA sequencing increased the gene and sample throughput while maintaining a degree of resolution of the tissue heterogeneity^8,12,13^. Additionally, the use of spatial transcriptomics has identified gradients of benign-cell gene transcription around tumour foci^14^. However, these studies mainly focused on the process of epithelial to mesenchymal transition^12,13^ (EMT) or were limited to the overall stromal contribution to disease progression^8^. An integrative investigation of immune, stromal and cancer cell transcriptional changes associated with clinical risk is still lacking.

In this study, we applied an optimised protocol for combined cell-type enrichment and ultra-low-input RNA sequencing, which allowed probing of four key cell types across 13 fresh prostate tissue spanning a wide clinical disease spectrum. Motivated by the pseudo-continuous properties of CAPRA-S risk score, we developed a novel statistical inference model for differential transcription analyses on continuous covariates, TABI (Transcriptional Analysis through Bayesian Inference). Our inference model was able to model changes in transcription without *a priori* patient risk stratification, and robustly map transcriptional change events to cancer risk states. Several hallmarks of prostate cancer were identified among the signatures of cell-surface and secreted protein coding genes. An emergent signature for monocyte-derived cell recruitment was identified and tested both with tissue deconvolution on the extensive Cancer Genome Atlas (TCGA) cohort and with multiplex immunohistochemistry on an independent patient cohort. The spatial analysis at the single-cell level allowed us to better identify relations between macrophages and epithelial and T cells along cancer progression.

## Results

### Data generation and quality

To investigate the role of the tumour microenvironment in patient outcome, we enriched for four cell populations (epithelial: EpCAM^+^; fibroblasts CD90^+^/CD31^−^; T cells: CD45^+^/CD3^+^; and myeloid: CD45^+^/CD16^+^) from fresh prostatectomies of 13 prostate cancers, ranging from benign tissue (labelled as CAPRA-S score 0) to high-risk tumours (CAPRA-S risk score 7). The choice of those cell populations was guided by their predominant role in the progression of prostate and other cancers^15–19^. Technical and practical experimental limitations prevented the consideration of other key cell types such as luminal and basal epithelial compartments, endothelial, smooth muscle and other lymphocytes such as B and natural killer cells. RNA extracted from the four cell populations was then sequenced, generating a median of 22 million reads per library (Fig. S1 and S2). Overall for the four cell type categories, a conservative cell-type purity inference (using Cibersort and a Bayesian estimator; Material and Methods) estimated high enrichment: 99% for epithelial samples; 99% for fibroblasts; 97% for myeloid (83% for neutrophils; 14% for monocyte derived cells); and 95% for T cells (Fig. S3). On average across the four cell types, 40% of genes had 0 sequenced reads in more than half of samples and were removed from further analysis.

### Differential transcription and model fitting

Dimensionality reduction (multidimensional scaling; MDS in Materials and Methods) of the filtered transcript abundance revealed an association of CAPRA-S risk score with either the first and/or the second principal components (Fig. 1A; with an indicative direction represented by the dashed grey line; tested with linear regression, lm function from R). A clear gradient in risk score was seen for epithelial and fibroblasts (Bonferroni adjusted p-value of 1.0×10^−2^ and 9.7×10^−3^ respectively), while a weaker pattern was apparent for myeloid and T cells (Bonferroni adjusted p-value of 3.0×10^−2^ and 0.37 respectively); possibly due to the greater heterogeneity of the two immune cell populations compared to epithelial and fibroblasts.

**Figure 1:**
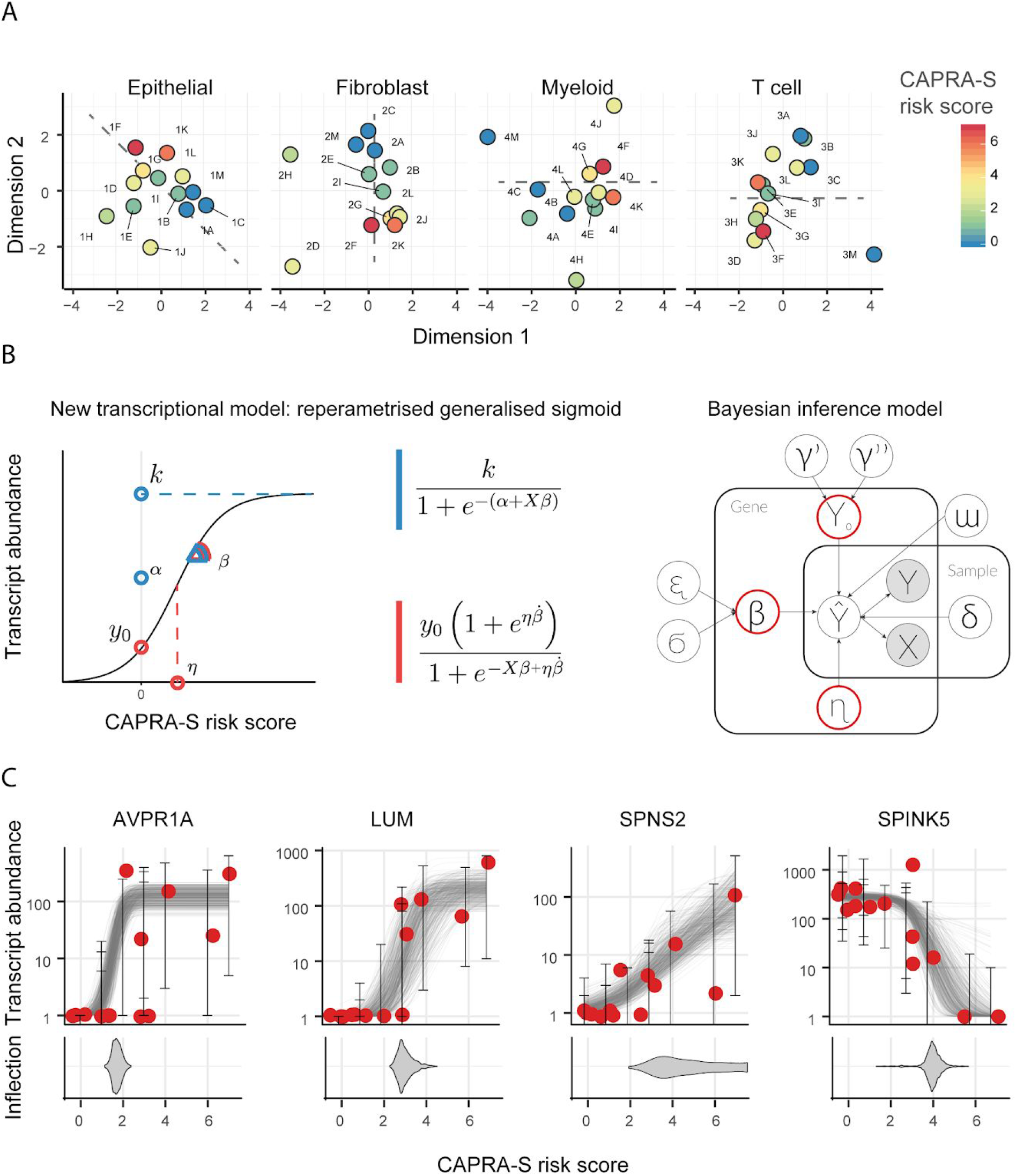
Data exploration and implementation of a model reflecting the continuous (i.e., non-discrete) relation between the CAPRA-S risk score and gene transcript abundance. **A** — Multidimensional scaling (MDS) plots of transcript abundance grouped by cell type. The colour coding represents the CAPRA-S risk score. The risk score shows a correlation (tested with linear regression, lm function from R) across the first and/or the second dimension (Bonferroni adjusted p-value of 0.0187, 0.00971, 0.0306 and 0.367, respectively), particularly in epithelial and fibroblast cells. Alphanumeric-codes refer to patient identifiers (Supplementary material). The dashed lines indicate the main direction of correlation between the first and/or the second dimension with CAPRA-S risk score. **B** — Representation of the re-parameterisation of the generalised sigmoid function and the resulting probabilistic model (Material and Methods). Left-panel: In blue are the three reference parameters for the standard parameterisation, in red are those for the alternative robust parameterisation. Right-panel: graphic representation of the probabilistic model TABI. **C** — Example for the continuous associations between transcript abundance of four representative genes and CAPRA-S risk score (for epithelial cell population), from more discrete-like to more linear-like. The bottom section displays the inferred distribution of possible values (as posterior distribution) of the inflection point for each gene sigmoid trend.

We performed a differential gene transcript abundance analysis for each cell type independently, seeking associations between transcript abundance across subjects and CAPRA-S risk score, treated as pseudo-continuous variables. In order to perform differential transcription analyses that would robustly model the pseudo-continuous properties of the CAPRA-S risk score (Fig. 1A), we developed a Bayesian inference model (TABI) that implements a robust generalised sigmoid regression (i.e., sigmoid function extending from zero to any positive value). TABI was used to model the gene transcript abundance as a continuous function of CAPRA-S risk score (from 0 representing benign to 7 representing high risk); this avoids the loss of information caused by the *a priori* stratification of patients in low-/high-risk groups based on an arbitrary threshold. In principle, the use of a generalised sigmoid function permits modelling linear, exponential and sigmoid-like trends of transcriptional alterations (Fig. 1B and 1C); however, to provide robust modelling for RNA sequencing data we reparameterized the generalised sigmoid function to better suit the numerical properties of transcript abundance (Fig. 1B; Materials and Methods). In addition to robustness, the sigmoid function allows the mapping of each differential transcriptional event with a clinical risk state, effectively providing a new developmental dimension to the analyses. This is possible because the inflection point represents the CAPRA-S risk score at which the transcriptional alteration is most pronounced. The location of most rapid change can be highly localised in the case of a dramatic change in transcription at a specific risk score or can be diffused in the case of a gradual change of transcription along the risk score range (Fig. 1C).

Following the statistical inference of transcriptional alterations, an average of 10% of genes were removed based on the posterior predictive check across all samples^20^, as not satisfying the assumptions of our model (Table 1; Materials and Methods). A total of 1,626 genes were identified as differentially transcribed across the four cell type categories (i.e., 95% credible interval excluding zero; with no need for multiple test adaptation, consistent with common practices in Bayesian statistics^21^; Table 1; Supplementary file 1). The distributions of differential transcription events along the CAPRA-S risk score range are concentrated on low risk scores (Fig. S4) for the four cell types, indicating that most transcriptional changes occur early in cancer developmental stages (including benign prostate tissue).

**Table 1:**
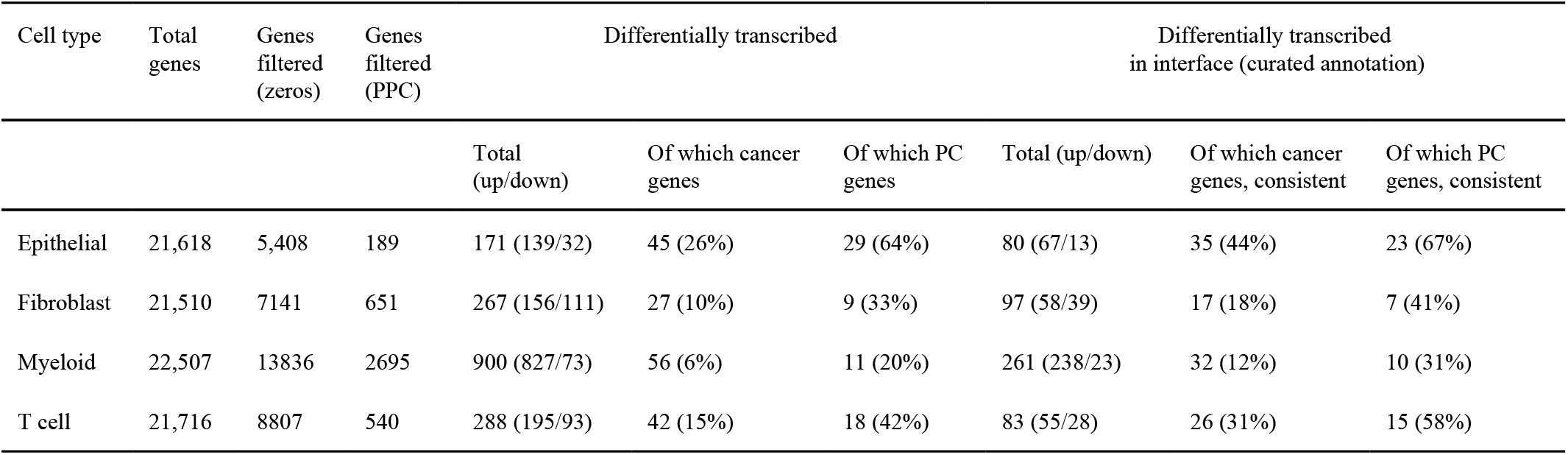
Summary statistics of the differential transcription analysis, including 52 samples from 13 patients and 4 enriched cell types. PPC = posterior predictive check; PC = prostate cancer. “Of which” refers to the gene selection relative to the category adjacent on the left. “Interface” refers to cell-surface and secreted protein coding genes. “Curated” refers to the curated database for cellular-interface genes produced in our study (Supplementary file 2). “Consistent” refers to a consistent direction of transcriptional change according to the curated database. Genes were labelled as “cancer genes” if present in the tier1 COSMIC database^95^ or labelled as such in our manually curated cell-type specific database (Supplementary file 2). Genes were labelled as “prostate cancer genes” if present in the tier1 COSMIC prostate cancer database dataset^95^ or labelled as such in our manually curated cell-type specific database (Supplementary file 2).

### Differentially transcribed cell-surface and secreted protein coding genes are linked with recurring cancer hallmarks

In order to provide an initial biological evaluation of the resultant differentially transcribed genes, we sought the overlap with cancer related gene datasets, and calculated the enrichment of gene sets against functional and clinical gene annotation databases. On average across the four cell types, 14% of all the differentially transcribed genes have been previously identified as cancer-related; of these, 24% have been previously described as prostate cancer-related genes (Table 1). For differentially transcribed cell-surface and secreted protein coding genes, an average of 33% and 51% have been previously described as cancer and prostate cancer-related genes respectively (Table 1). In order to investigate possible cell-cell interactions within the primary prostate tumour microenvironment, we focused on genes encoding for cell-surface and secreted proteins, which may have direct influence on other cell types. On average across the four cell types, 35% of differentially transcribed genes encode for cellular-interface proteins; of those, 148 genes have been previously described as cancer related genes. For all cell types, most cancer genes have a direction of change consistent with the direction reported in the literature (35 vs. 13 for epithelial; 17 vs. 8 for fibroblasts; 32 vs. 6 for myeloid cells; and 26 vs. 11 for T cells; Supplementary file 3).

In order to allow an in depth interpretation of the concurrent transcriptional differences for cell-surface and secreted protein coding genes across cell-types, we produced a cell-type and disease specific annotation database integrating curated cell-specific Gene Ontology information^22^ with more than 1500 scientific articles (Supplementary file 3). This allowed us to identify six recurring hallmarks of cancer (Fig. 2): (i) immune modulation; (ii) cancer cell migration; (iii) angiogenesis; (iv) hormonal homeostasis; (v) epithelial/cancer cell growth; and (vi) osteogenesis. Among the immune modulation related genes, a balance exists between pro- and anti-inflammatory. This balance appears to be dynamic along the disease progression course. The epithelial cell migration hallmark includes three main functional clusters: tissue remodelling, tissue fibrosis and direct epithelial-to-mesenchymal transition. The differential transcription events of those three classes do not appear to be concentrated on any particular stage of disease progression. Similarly, angiogenesis signalling appears to be sustained along the whole disease progression, where a gene alteration signature of platelet recruitment linked with endothelial cell migration is expressed in synergy by both myeloid and T cells. Several transcriptional alterations from both epithelial and immune cells were linked with hormonal and lipid homeostasis, which is a key molecular hallmark in prostate cancer^23^. Within this set, the most recurring metabolite that is linked with differentially transcribed genes is cholesterol. While for most hallmarks all four cell types contributed similarly to the signatures, a clear bias is present for cancer cell growth, osteogenesis and hormone modulation signatures, which are enriched with genes that are differentially transcribed in epithelial and immune cells types respectively. As the most compelling signal, immune modulation was selected for further investigation.

**Figure 2.**
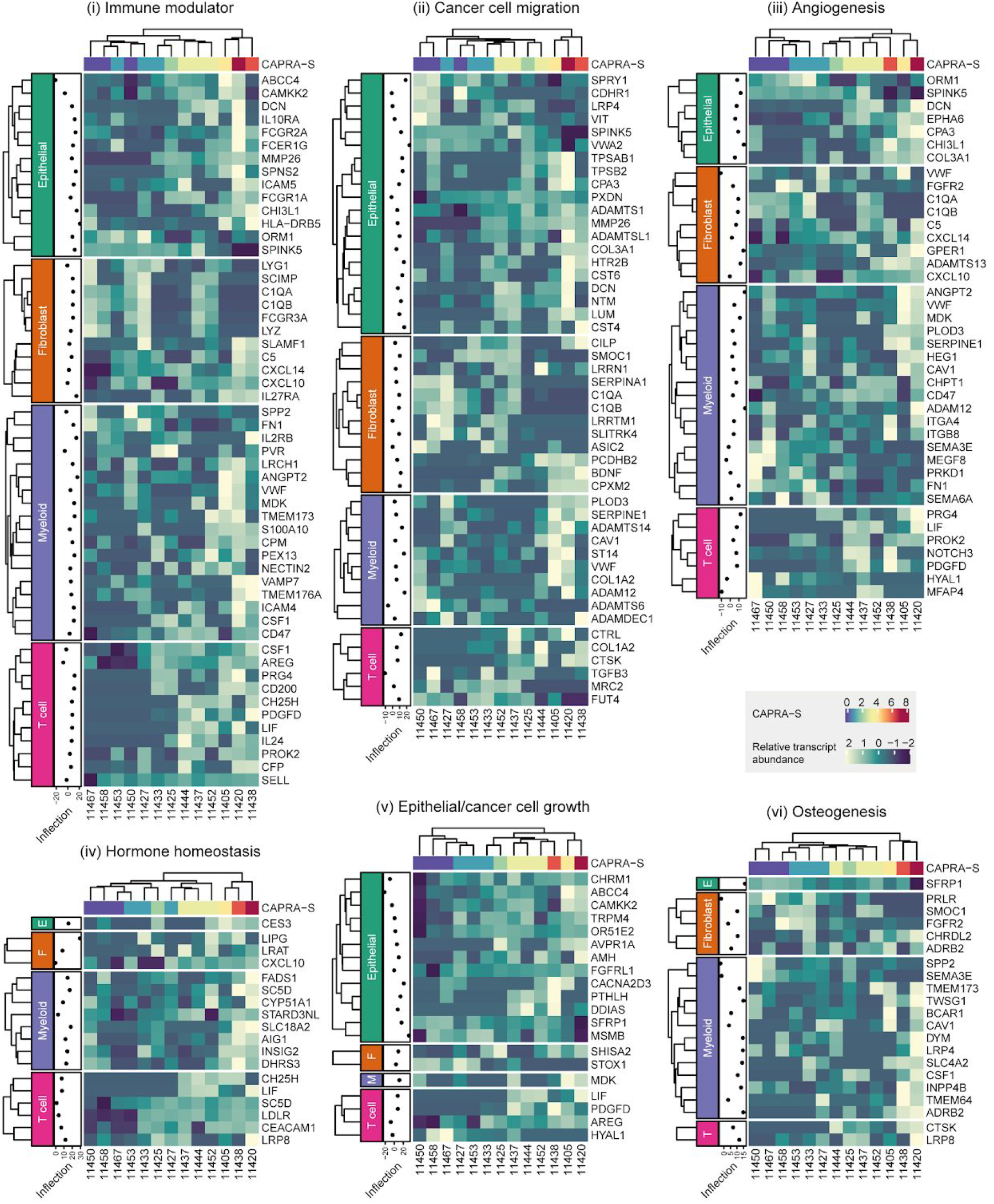
Recurrent functional categories identified in the differentially transcribed cellular interface-protein-coding (secreted and transmembrane) genes. The estimated inflection point for each gene shows the CAPRA-S risk score at which the transcriptional change was inferred to be fastest; values < 0 or > 7 indicate an early or late change, respectively.

### Immune modulation is associated with cancer grade and targets predominantly monocyte-derived cells

In order to elucidate the role of the four cell types in the immune response to primary prostate cancer and their potential interactions, we focused on genes that encode for cell-surface and secretory proteins involved in immune modulation. In doing so, we again used the fitted inflection point of the sigmoid model to distinguish between early (i.e. low CAPRA-S risk score) and late (i.e. high) transcriptional changes. The balance between pro- and anti-inflammatory signatures from the four cell types tracks with the risk score covariate (Fig. 3). While the magnitude of the pro-inflammatory transcriptional signature remains roughly constant through the risk range, with 18 genes for CAPRA-S risk score ≤ 2 and 14 for CAPRA-S risk score >2; the anti-inflammatory signature significantly expands (p-value 0.015; t-test) for more advanced stages of the disease, with 12 genes against 20 for the two risk score categories respectively.

**Figure 3:**
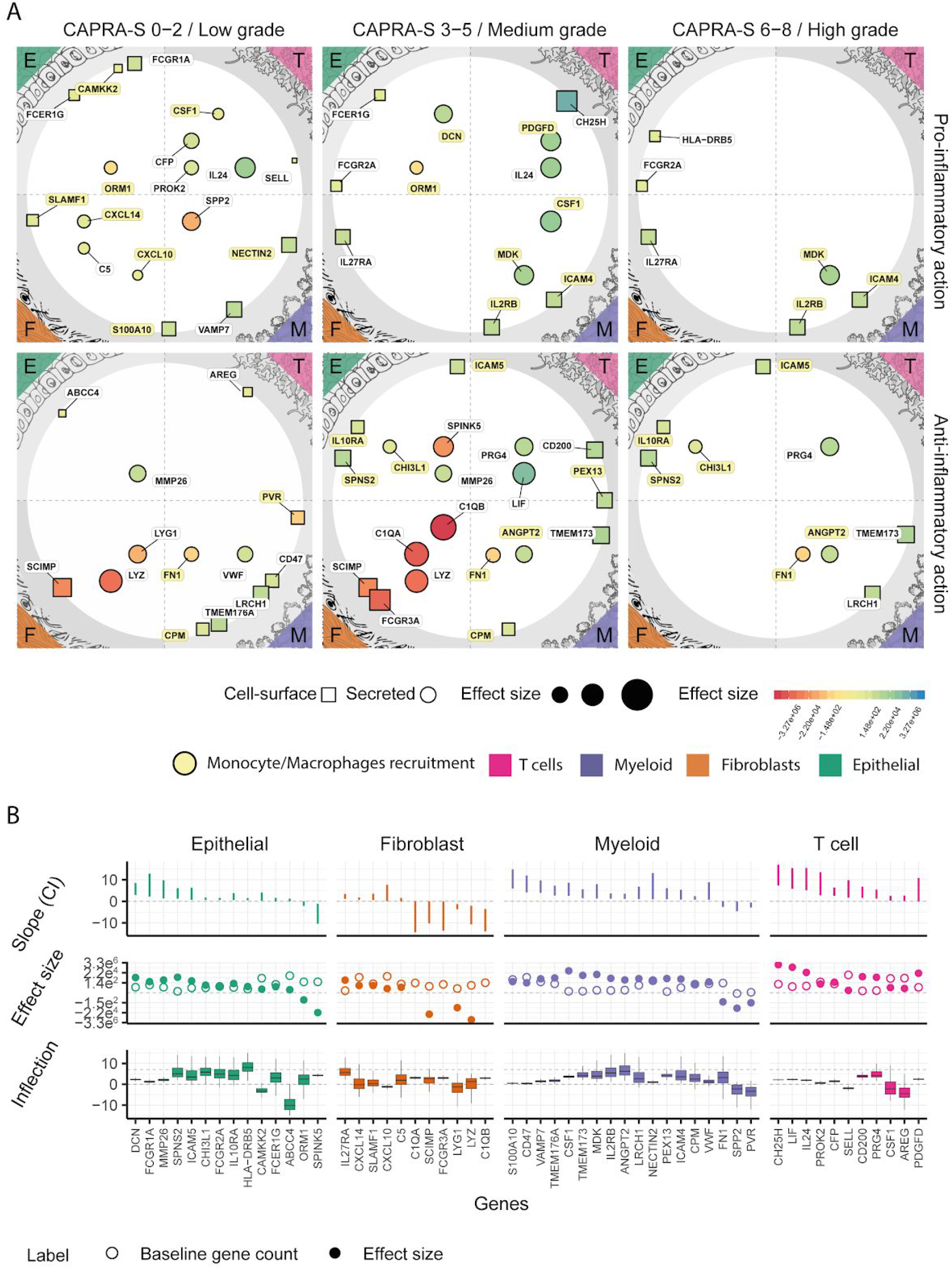
Multi cell-type immune-modulation evolves with risk progression and is mainly targeted at monocyte-derived cells. Landscape of the immune-modulation related genes that encode for cellular interface-proteins (i.e., cell-surface or secreted) inferred by TABI to be differentially transcribed across CAPRA-S risk scores, grouped by cell type. **A** — Map of the secretory (represented as circles) and cell-surface (represented as squares) protein-coding genes, that are differentially transcribed across the four cell types. The data point size is proportional with the baseline transcript abundance. The colour coding represents the effect size. Genes with a similar inflection point (i.e., at what stage of the disease a transcriptional change happens) are clustered vertically (CAPRA-S risk score <=2, >2 and <=5 and >5). Genes are split horizontally according to their pro- or anti-inflammatory role. Genes encoding for proteins that target monocyte-derived cells are highlighted in yellow. **B** — Statistics of the differentially transcribed genes displayed in panel A. **Top** — credible interval of the association between transcript abundance and CAPRA-S risk score. **Middle** — inferred effect size (full dots) and baseline transcription (empty dots). **Bottom** — credible interval of the CAPRA-S value for the transcriptomic change (i.e., inflection point; e.g., the gene HLA−DRB5 is upregulated in late stages of the disease).

Overall, a large proportion (14 genes of 27) of the inflammatory-related transcriptional alterations across all four cell types is involved in the recruitment of monocytes and macrophages^24–31^ (highlighted in yellow in Fig. 3A). These include CAMKK2^32^, ORM1^33^ and DCN^34,35^ in epithelial; IL2RB^36–38^, ICAM4^39,40^, DCN^34,35^ and MDK^41,42^ in myeloid cells; and CSF1^24^ and PDGFD^43^ in T cells. In addition, we identified a known fibroblast-macrophage chemotactic interaction including the regulation of the cytokines CXCL10^27^, CXCL14^26^ and the receptor SLAMF1^28,29^ for fibroblasts; with COL1A2^44^ and CYR61^45^ (for CAPRA-S 6-8) altered in myeloid cells known to function as co-stimulatory loop. A smaller cluster of genes were linked with T cell recruitment and inflammation, including CFP^46^, IL24^47^, PROK2^48^, SELL^49^. Interestingly, epithelial cells upregulate a cluster of receptor genes normally involved in antigen recognition and presentation in immune cells^50^, including an MHC class II cell surface receptor (i.e., HLA-DRB5) and three Fc receptors (i.e., FCER1G, FCGR1A and FCGR2A).

The anti-inflammatory signatures target a more heterogeneous set of cell types than the pro-inflammatory signature. Monocyte-derived cells are mainly targeted by genes that are differentially transcribed in epithelial and myeloid cells. These include the receptor genes SPNS2^51,52^, IL10RA^53^ and ICAM5 by epithelial cells; and the receptor genes CPM^54^ and PEX13^55^ and the secreted protein genes FN1^56^ and ANGPT2^57^ by myeloid cells. Another cluster of genes targets predominantly T cells, including AREG^58,59^, CD200^60^, LRCH1^61^, CD47^62^. Fibroblasts mainly downregulate pro-inflammatory cell-surface and secreted protein genes, such as FCGR3A and C1QA/B.

### Increased monocyte-derived cell infiltration in tumours is associated with lowered disease-free survival

In order to test the relevance of recruitment of monocyte-derived cells suggested by our integrated transcriptional analysis, we performed a differential tissue composition (DTC) analysis (i.e. a test for difference in cell-type abundance between conditions) based on an independent methodology and independent patient cohort. We used a hierarchical Bayesian inference model^63^ on an independent cohort of 134 patients from the primary prostate cancer TCGA dataset^7^ that included both disease-free survival and CAPRA-S score information. This algorithm uses a collection of 250 curated publicly available transcriptional profiles (including BLUEPRINT^64^, ENCODE^65^, GSE89442^66^ and GSE107011^67^), encompassing transcriptional signatures for 18 hierarchical cell-types (of which nine major cell types were tested for differences) and uses those reference signatures to understand the contribution of each cell type to the observed mixed transcriptional signal. This analysis provides tissue composition estimates as well as their association with risk score^63^. Overall, we estimated a median of 88% for epithelial cellular fraction across samples (consistent with public literature^68^, Fig. S5), 4.8% for endothelial, 4.8% for fibroblasts, and 1.6% for immune cells. The differential tissue composition analysis showed a significant positive association with the CAPRA-S risk score of the monocyte-derived, and a negative association of the natural killer and granulocyte cells (95% credible interval excluding 0; Fig. 4A).

**Figure 4.**
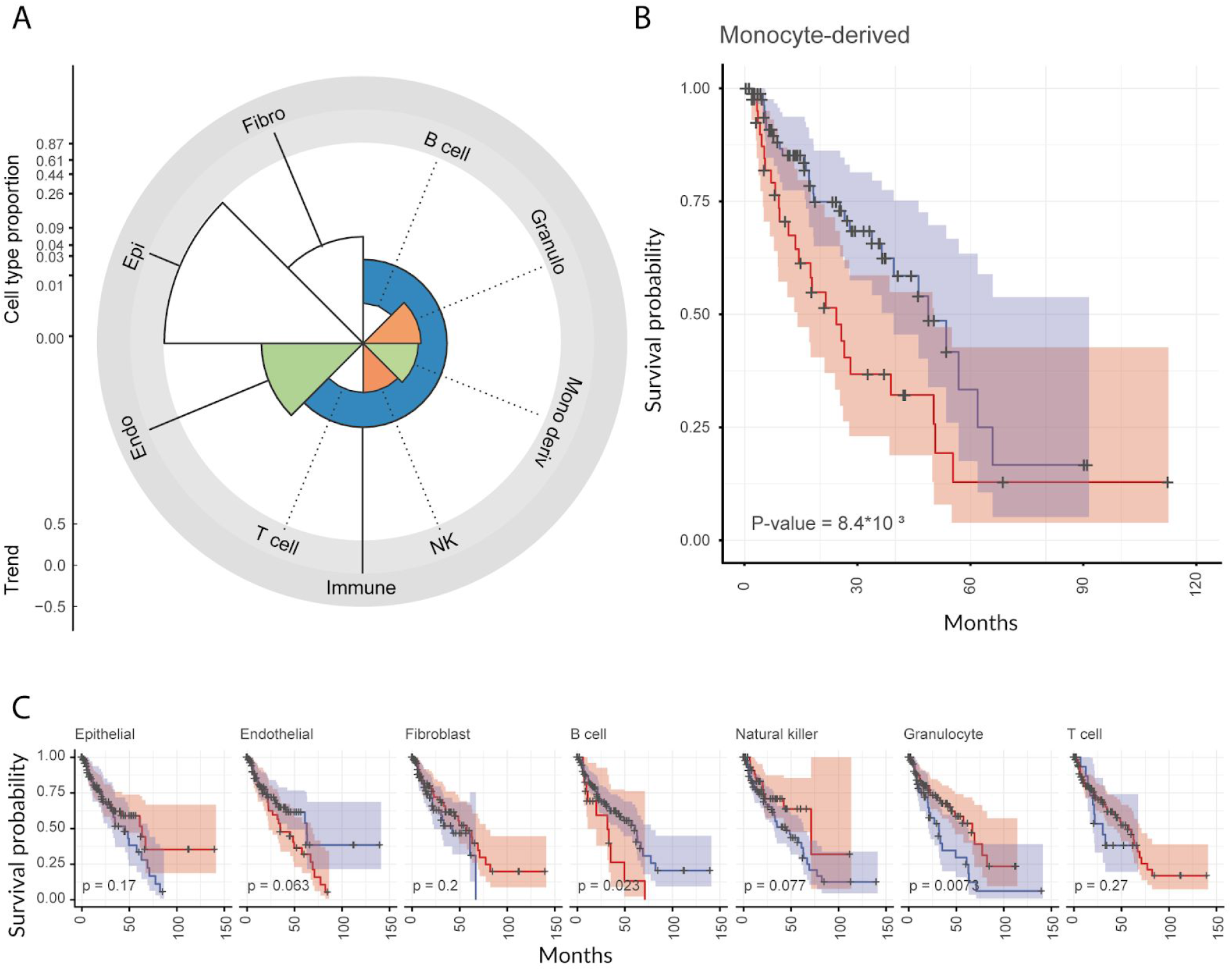
The abundance of monocyte-derived cells is positively associated with CAPRA-S risk score and negatively associated with disease-free survival. Association analysis of the abundance of monocyte-derived cells (inferred with a Bayesian model) with disease-free survival, performed on the independent primary prostate cancer TCGA dataset (n = 134). **A** — Polar plot for the differential tissue composition analysis of primary prostate cancer TCGA samples (n=134) for which CAPRA-S risk score information is available, with the factor of interest being CAPRA-S risk score. The y-axis (scaled by the fourth root) represents the overall cell type abundance; the colour coding reflects the association between cell type abundance and disease-free survival (coloured = significant association). **B** — Kaplan–Meier plot of patients (n = 134) with low (blue) or high (red) monocyte-derived cell infiltration in the tumour specimen (proportion cut-off = 0.0048; see Material and Methods section, Survival analyses subsection). **C** — Kaplan–Meier plot for the other cell types included in the analysis.

In order to test whether the enrichment in monocyte-derived cells is clinically relevant, we generated Kaplan–Meier curves using the estimated cell-type abundances. The stratification of patients based on the extent of monocyte-derived cells infiltration revealed a significant separation of the survival probabilities (Fig. 4B). For comparative purposes, we tested patient stratification for the other cell types included in the model. As a result, only granulocytes and B cells (with the poor outcome cohort including only few patients) showed a significant negative association, while no other significant associations were detected for other cell types including epithelial, endothelial, fibroblasts, and immune cell types such as including T cells and natural killers (Fig. 4C).

### Macrophage proximity to epithelial glandular clusters increases with tumour progression

In order to validate our findings and gather more in depth knowledge about the role of macrophages in disease progression, we performed a spatial analysis at the single-cell level of 63 prostate biopsies from the independent RadBank cohort of 17 patients with localised PC - spanning a wide range of CAPRA risk scores (Table S1). Using 7 immuno-fluorescent dyes (including DAPI) with 6 others linked to antibodies (CD3, CD20, CD68, CD11c and HMWCK), cell size and shape we were able to identify 6 major cell types: T-cells, B-cells, macrophages, dendritic myeloid, epithelial basal and stromal cells. PDL1 expressing cells were also categorised, using a PDL1-linked dye. Overall, we were able to sample an average of 41×103 cells per biopsy (standard deviation 2.7×10^4^). Of the categorised cell types, the most abundant was epithelial basal (8.06% on average), followed by T-cells (5.03% on average).

We estimated the association between macrophage proximity to five other cell types and CAPRA risk score (Fig. 5A). Across the five cell types, the average number of neighbour cells to macrophages ranges from 1.4 to 23.0. The strongest positive association was between macrophage and epithelial basal cells in tumour areas (p-value 0.031; Fig. 5A-top-right), while the strongest negative association was for stromal cells in tumour areas (p-value 0.093; Fig. 5A-bottom). Overall, the average proximity between macrophages and other cells, aggregated by biopsy did not show any strong association with CAPRA risk score. In order to gather evidence that the increased proximity of macrophages to the prostatic gland structures in advanced cancer stages had some direct effect on the local immune microenvironment, we tested the hypothesis that macrophages in proximity to gland structures would displace T-cells. We observed an inverse association between the number of neighbour macrophage expressing PDL1 to epithelial basal cells, and the number of T-cell neighbours (Fig. 5B). That is, epithelial basal cells that are close to clusters of PDL1 expressing macrophages tend to be further away from T-cells.

**Figure 5.**
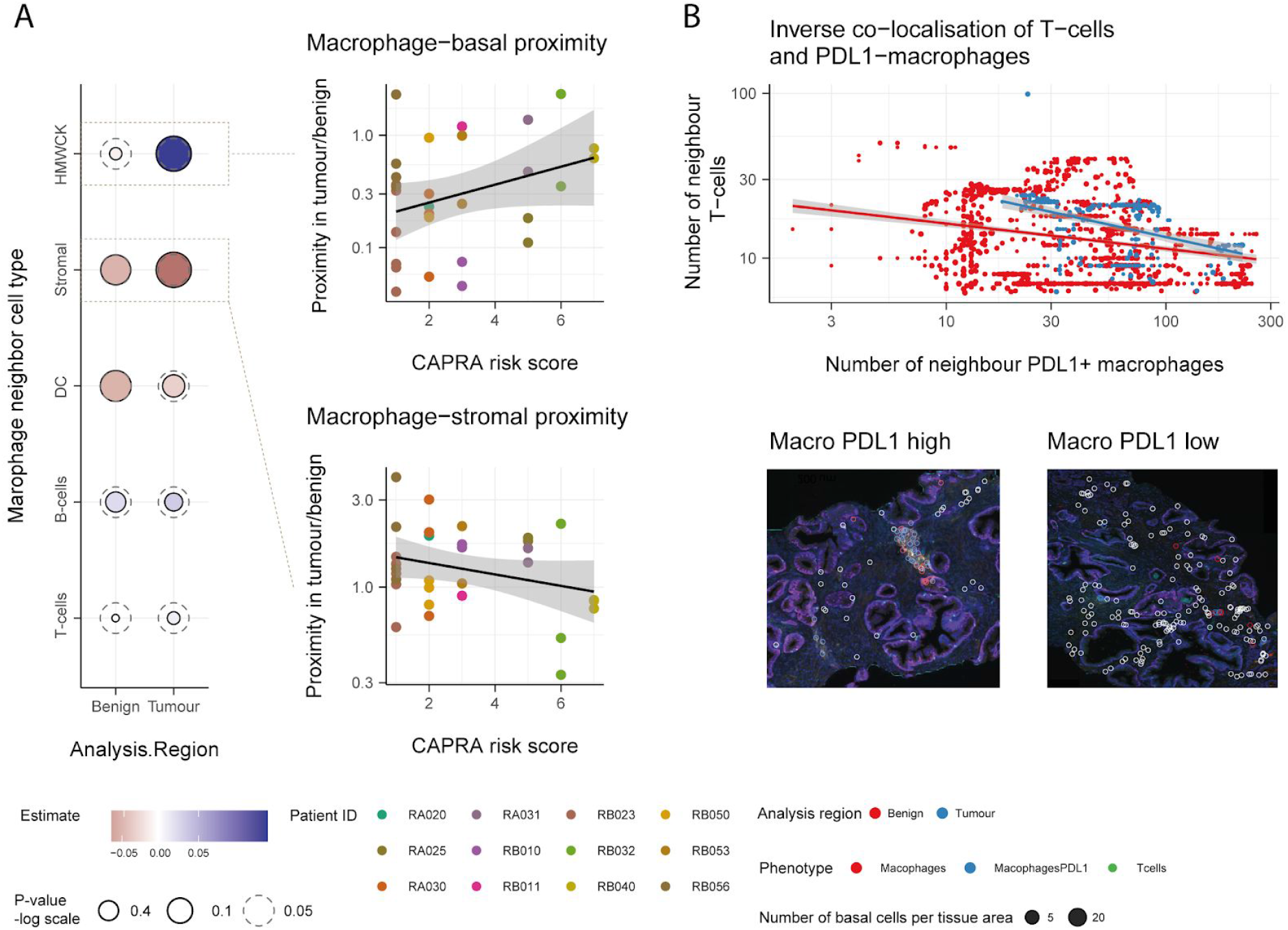
**A left** - Association between macrophage proximity and CAPRA risk score for five cell types identified from the multiplex immunohistochemistry. Proximity is calculated as number of neighbour cells per tissue area, and summarised using the median for each tumour biopsy. **A right** - Association between macrophages and epithelial basal cells (top) or stromal cells (bottom) and CAPRA risk score shown in panel A. Only the 12 patients with both tumour and surrounding benign tissue are displayed. **B** - Decreased proximity of T-cells with epithelial basal in presence of PDL1 expressing macrophages. The bottom section shows the multiplex immunohistochemistry tissue from patient RB010, with two examples of presence (left) or absence (right) of PDL1 macrophages close to prostate glandulae. White circles surround the labelled T cells, blue and red circles surround macrophages which are PDL1 low and high respectively.

## Discussion

To date, in-depth analyses of genomic features of prostate cancer alone, including single nucleotide variants and small and large structural rearrangements, have not been sufficient to provide transformative prognostic tools or unveil the full complexity of this disease. It is clear that non-malignant cells within the tumour microenvironment make an integral contribution to the mechanisms that cause cancer progression, and that they are often modulated by cancer cells toward pro-tumorigenic behaviours. In this study, we significantly improved the knowledge of the molecular landscape of primary prostate tumour microenvironment, revealing concurrent transcriptional changes in epithelial, fibroblast, myeloid and T cells along the CAPRA-S risk score range.

We optimised a combined fluorescence-activated cell sorting and ultra-low-input RNA sequencing protocol, allowing us to obtain high quality sequencing data from inputs down to 1000 cells. Such a strategy is of general utility as it enables studies of rare cell types from both fresh tissue cores and biopsies. In order to optimally detect changes in transcription along CAPRA-S risk score, we developed a novel statistical inference method (TABI), which permitted modelling of transcript abundance natively on continuous factors of interest with a limited number of parameters, avoiding loss of information due to the dichotomization of the risk score into low-/high-risk patient groups. As suggested by multidimensional-scaling plots and supported by our inference, transcriptional change events are indeed continuously distributed along the whole risk score range. This method is of broad utility in all cases where a continuous (or pseudo-continuous) factor of interest is present (e.g., risk score, time and chemical concentration) and a monotonic change in transcript abundance is of interest. Furthermore, the novel parametrisation of the generalised sigmoid function that TABI is based on, can be extrapolated for a wide range of applications.

The compilation of a curated cell-type specific database of gene functions for cell-surface and secreted protein coding transcripts enabled the detection of several recurrent hallmarks of prostate cancer, characterised by the involvement of multiple cell types. The most striking aspect to emerge was the large number of differentially transcribed genes linked to monocyte-derived cells recruitment and modulation. The association of monocyte-derived cell recruitment with increased risk score was reflected in an orthogonal differential tissue composition analysis on the extensive Cancer Genome Atlas (TCGA) independent patient cohort against CAPRA-S risk score, through a differential tissue composition analysis. This analysis was enabled by a robust Bayesian inference model, able to transfer the uncertainty of the estimation of tissue composition for each sample to the linear model linking cell-type proportion to clinical variables. Such an aspect is particularly relevant considering the substantial noise associated with such inference. To test the clinical significance of quantifying monocyte-derived cell numbers within the tumour mass beside the CAPRA-S risk score, we performed survival analyses from the same TCGA cohort, from the inferred cell-type proportions. For both analyses, we identified the strongest association with clinical variables being monocyte derived cells. The infiltration of cell types such as monocyte-derived cell populations has previously been shown to be linked to the extent of proliferative inflammatory atrophy lesions, chronic prostatic inflammation and cancer grade^69^. In prostate cancer, specific and overall survival analyses have identified an elevated monocyte count as an independent prognostic factor for poor outcome^70–73^. Furthermore, the infiltration of tumour associated macrophages in prostate needle biopsy specimens has been shown to have potential as a predictive factor for PSA failure or disease progression after hormonal therapy^74^.

In order to validate further our hypothesis and enrich our knowledge about the relation of macrophages with epithelial and a range of immune cells along disease progression, we used multiplex immune-histochemistry (IHC) to determine the immune context at the single cell level in an independent cohort. This data supports the hypothesis of a weakened relation of macrophages with stromal compartments and a strengthened association with epithelial glandular clusters along the disease progression spectrum, while distanciating from stromal compartments; the latter being colocalized with cancer cells in tumour tissues This aspect becomes highly relevant as the glandular areas in both tumour and adjacent benign compartments that are rich in PDL1 macrophages become consequently poorer in T-cell numbers. PDL1 expressing macrophages have been associated with their M2 wound-healing phenotype^75^. The relationship between PD-L1 expression in intratumoral macrophages and prognosis in cancer patients is still controversial, with the two competing hypotheses being: (i) PDL1 intratumoral macrophages lead to dysfunctional T-cells^76^ or (ii) not having major effects^77^. Nonetheless, expression of PDL1 in macrophages has been shown to induce anti-inflammatory cytokines such as IL-1075. Although PDL1 in macrophages may primarily function as protection against induced cell death, our study supports the hypothesis that it may have the effect of inducing an anti-inflammatory, immune cold local microenvironment, with negative effects on disease progression.

## Conclusions

In prostate cancer, there has been limited benefit observed through the unselected use of novel immune checkpoint inhibitors based on T cell receptor blockade (e.g., PD-1, PD-L1 and CTLA-4)^78^. Such failure may in part be driven by our limited understanding of the dynamic interplay between immune components of the microenvironment and tumour cells. This study provides a clear direction for further investigation into mechanisms of the immune system (particularly, monocyte-derived cells) that contribute to disease progression; for example, changing the hormonal and growth-factor homeostasis through a sustained inflammatory state. Furthermore, this study provides a novel and robust method for detecting monotonic changes in transcript abundance over a continuous factor of interest such as risk and time, that has broad applicability to other research areas. Both the methodological advances and the novel findings presented in this study provide a research framework for improved immune interventions.

## Declarations

### Ethics approval and Consent to participate

The collection and use of tissue for this study had Melbourne Health institutional review board approval and patients provided written informed consent (Melbourne Health Local Project Number: 2016.087; and PMCC; HREC approvals 10/68, 13/167, 18/204).

### Availability of data and materials

The code used to conduct the analyses is available at github.com/stemangiola under the following repositories: prostate-TME-N52-2019; TABI@v0.1.3; ARMET@v0.7.1. Sequence data has been deposited at the European Genome-phenome Archive (EGA), which is hosted by the EBI and the CRG, under accession number EGAD00001004948.

## Acknowledgements

We wish to thank Dr. Stephen Wilcox for technical assistance and are grateful to the staff of the Flow cytometry facility and Clinical Translational Centre (Walter and Eliza Hall Institute of Medical Research). We thank Professor Risbridger (Monash University) for the great support with the initial phase of the study. We thank Dr. Damiano Spina (RMIT) for the support with machine learning. We thank Dr. Bob Carpenter and Prof. Andrew Gelman (Columbia University) for an incredible mentorship in Bayesian inference, and all the Stan community at discourse.mc-stan.org for technical support. Most importantly, we thank all the patients who participated in the study. We thank the anonymous reviewers whose comments have greatly improved this manuscript.

## Authors’ contribution

SM conceived and designed the study, performed part of the cellular biology procedures, implemented the statistical methods and performed data analysis and visualisation, under the supervision of AJC, NH, NMC, ATP and CMH. MK, JP and PD contributed with sample harvesting. PM, FSFG, DB, NL, CN, MLP and SIM contributed to the cellular biology procedures; and PM and BP contributed to the molecular procedures. MM contributed to statistical model implementation. SPK led the multiplex histochemistry experiments supervised by PJN and SGW. All authors contributed in manuscript writing.

## Competing interests

The authors declare that there is no conflict of interest that could be perceived as prejudicing the impartiality of the research reported.

## Funding

SM was supported by the David Mayor PhD Scholarship from the Prostate Cancer Research Foundation, and by the Galli Next Generation Cancer Discoveries Initiative. KC was supported by a Postgraduate Medical Research Scholarship from the Prostate Cancer Research Fund, and the Australian Government Research Training Program Scholarship provided by the Australian Commonwealth Government and the University of Melbourne. NDH was a recipient of a Melanoma Research Grant from the Harry J Lloyd Charitable Trust. FSFG was supported by a grant #1158085 awarded through the Priority-driven Collaborative Cancer Research Scheme and funded by Cure Cancer Australia with the assistance of Cancer Australia. MM was supported by ELIXIR CZ research infrastructure project (MEYS Grant No: LM2015047) including access to computing and storage facilities.. NMC was supported by a David Bickart Clinician Research Fellowship from the Faculty of Medicine, Dentistry and Health Sciences at the University of Melbourne, as well as Movember — Distinguished Gentleman’s Ride Clinician Scientist Award through Prostate Cancer Foundation of Australia’s Research Programme. ATP was supported by the Lorenzo and Pamela Galli Charitable Trust and by an Australian National Health and Medical Research Council (NHMRC) Program Grant (1054618) and NHMRC Senior Research Fellowship (1116955). The research benefitted by support from the Victorian State Government Operational Infrastructure Support and Australian Government NHMRC Independent Research Institute Infrastructure Support.

## Material and methods

### Tissue sampling and processing

Following the prostatectomy of 13 patients, ranging from 52 to 78 years of age and from CAPRA-S risk score of 0 (attributed to benign tissue samples, harvested from a site far from a low grade, low volume cancer) to 7 (Supplementary file 4), a four millimeter tissue core was collected from the prostate tumour site, conditional to histopathological verification^79,80^. If not otherwise specified, all procedures were carried out at 4 °C. Tissue blocks were washed in Phosphate-buffered saline (PBS) solution for 2 minutes and minced for 2 minutes with a scalpel. Homogenised tissue was added to a solution (total volume of 7 ml) composed by of 1 mg/ml collagenase IV (Worthington Biochemical Corp, USA), 0.02 mg/ml DNase 1 (New England Biolabs, USA), 0.2 mg/ml dispase (Merck, USA). The homogenised tissue was serially digested in the shaker incubator at 37 °C at 180 rpm (4g), through three steps of 5, 10 and 10 minutes of duration, with the final 3 minutes dedicated to sedimentation at 0 rpm. After each digestion step, the supernatant was aspirated and filtered through a 70 μm strainer into a pre-chilled tube, diluting the solution with 15 ml of Dulbecco’s PBS containing 2% Bovine serum (dPBS-serum) to quench the enzymatic reaction. The resulting cumulative solution was then centrifuged at 300gfor five minutes, with the supernatant collected and the cell pellet resuspended into 1 ml 2% PBS-serum prior to labelling (Fig. S1).

### Antibody labelling, flow cytometry and cell storage

The cell preparation was labelled with the following antibodies: CD3-BV711 (Becton Dickinson San Jose Ca), EpCAM-PE (BD Biosciences, USA), CD31-APC (BD Biosciences, USA), CD90-PerCP-Cy5.5 (Becton Dickinson San Jose Ca), CD45 APC-CY7, and CD16 Pacific Blue (BD Biosciences, USA), Thy-1 (CD90) PerCP-Cy5.5 (BD Biosciences, USA). All antibodies were used at concentrations according to manufacturers recommendations and incubated for 30 mins at 4°C. Following labelling the cells were diluted to 5ml and centrifuged at 300g for 5 minutes. The supernatant was removed and the cell pellet was resuspended in dPBS-serum. Viability die (7AAD) was added to the suspension to a final concentration of 5μg/ml. Epithelial, fibroblasts, myeloid and T cells were sorted using a FACS Aria III cell sorter (Becton Dickinson San Jose, Ca). The cell sorting strategy utilized a robust 3 stage design: (i) a series of gates based or forward and side scatter to exclude debris, cell clumps and doublets. (ii) a gate to exclude all dead cells and (iii) combination of the fluorescent antibodies to allow purification of the above cell types. The four cell types were identified as follows: T-Cells: FSC and SSC lo, PI negative, EpCAM and CD31 negative, CD3 and CD45 positive. Epithelial cells: FSC and SSC high, PI negative, CD31 and CD90 negative and EpCAM positive. Myeloid cells: FSC and SSC hi and medium, PI negative, CD31 and EpCAM negative and CD16 positive. Fibroblasts: FSC and SSC hi, PI negative, EpCAM and CD31 negative, CD90 positive. The four purified populations were sorted directly into 1.5ml conical tubes and stored on dry ice immediately after collection before permanent storage at −80 °C.

### RNA extraction, library preparation and RNA sequencing

RNA extraction was performed in two batches (comprising 6 and 7 patients, for a total of 24 and 28 samples respectively) on consecutive days. In order to eliminate time-dependent methodological biases, the two patient batches included a balanced distribution of Gleason score (means 2.00 and 2.71, standard deviations 2.50, 1.86; Supplementary file 4) and days elapsed from tissue processing (means 197 and 222, standard deviations 46.3 and 71.9; Supplementary file 4). The RNA extraction was performed using the miRNeasy Micro Kit (Qiagen; Cat #217084), according to the manufacturer’s protocol. Briefly, cell pellets were lysed with QIAzol lysis reagent, treated with chloroform and centrifugation carried out to separate the aqueous phase. Total RNA was precipitated from aqueous phase using absolute ethanol, filtered through the MinElute spin column and treated with DNase I to remove genomic DNA. The RNA bound columns were washed with the buffers RWT and RPE before eluting the total RNA with 14μl of RNase-free water. RNA estimation was carried out using Tapestation (Agilent).

Transcriptome sequencing on low input total RNA samples (up to 10 ng) was carried out using SMART-Seq v4 Ultra Low Input RNA Kit (Clontech), according to manufacturer’s protocol. The first-strand cDNA synthesis utilised 3’ SMART-Seq CDS Primer II A and the SMART-Seq v4 Oligonucleotide together with the cDNA amplification was carried out on Thermocycler using PCR Primer II A and PCR conditions: 95 °C for 1 minute, 12 cycles of 98 °C 10 seconds, 65 °C 30 seconds and 68 °C 3 minutes; 72 °C for 10 minutes and 4 °C until completion. The PCR-amplified cDNA was purified using AMPure XP beads and processed with the Nextera XT DNA Library Preparation Kits (Illumina, Cat. # FC-131-1024 and FC-131-1096) as per the protocol provided by the manufacturer.

Sequencing library preparation (10 – 100 ng) was carried out using Truseq RNA Sample Preparation Kit v2. The poly-A containing mRNA was purified using oligo-dT bound magnetic beads followed by fragmentation. The first strand cDNA synthesis utilised random primers and second strand cDNA synthesis was carried out using DNA Polymerase I. The cDNA fragments then underwent an end-repair process, the addition of a single ‘A’ base, and ligation of the RNA adapters. The adaptor ligated cDNA samples were bead-purified and enriched with PCR (15 cycles) to generate the final RNAseq library.

The SMART-Seq v4 RNA and Truseq RNA libraries were sequenced on an Illumina Nextseq 500 to generate 15-20 million 75 base pairs paired-end reads for each sample. The batch effect due to sequencing runs was minimised by pooling all 52 libraries and carrying out three sequential runs on a Nextseq500 sequencer.

### Sequencing data quality control, mapping and read counting

The quality of the sequenced reads for each sample was checked using the Fastqc^81^. Reads were trimmed for custom Nextera Illumina adapters; low quality fragments and short reads were filtered out from the pools using BBDuk (jgi.doe.gov) according to default settings. All remaining reads were aligned to the reference genome hg38 using the STAR aligner^82^ with default settings. The quality control on the alignment was performed with RNA-SeQC^83^. For each sample, the gene transcription abundance was quantified in terms of nucleotide reads per gene (read-count) using FeatureCounts^84^ with the following settings: isPairedEnd = T, requireBothEndsMapped = T, checkFragLength = F, useMetaFeatures = T. All sequenced reads that did not align to the reference human genome were assigned to bacterial and viral reference genomes using kraken^85^ with default settings.

### Statistical inference

Changes of transcriptional levels along CAPRA-S risk score^86^ were estimated independently for each cell type (epithelial, fibroblast, myeloid and T cell). The CAPRA-S risk score is a combination of: (i) concentration of blood prostate serum antigen (PSA); (ii) presence of surgical margin (SM); (iii) Gleason score; (iv) presence of seminal vesicle invasion (SVI); (v) the extent of extracapsular extension (ECE); and (vi) lymph node involvement. The RNA extraction batch was used as further covariate. Due to the absence of publicly available models for non-linear monotonic regression along a continuous covariate, a new Bayesian inference model was implemented. This model is based on the simplified Richard’s curve^87^ (Eq.1), but re-parameterised to improve numerical stability (Eq. 2). In particular, the standard parametrisation suffers from non-determinability issues in case the slope is close to zero; furthermore, in case of an exponential-like trend the upper plateau is not supported by data and tends to infinity.

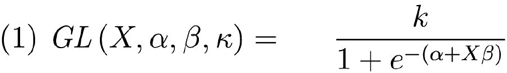

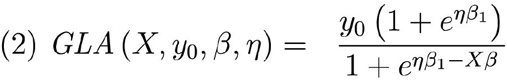

The new parameter y_0_ represents the intercept on the y axis, η represents the point of inflection on the x-axis, β represents the matrix of coefficients (i.e., slope coefficients, without the intercept term), β_1_ represents the coefficient of interest (i.e., main slope), and k the upper plateau of the generalised sigmoid function.

Bayesian inference was used to infer the values of all parameters of the model, with TABI (GitHub: stemangiola/TABI@v0.1.3). The probabilistic framework Stan^88^ was used to encode the joint probability function of the model (Eq. 3), partitioning the transcriptomic dataset in blocks of 5000 genes to decrease the analysis run-time. This Bayes model is based on a negative binomial distribution (parameterized as mean and overdispersion) of the raw transcript abundance. In order to account for diverse sequencing depths across samples a sample-wise normalisation parameter was added to the negative binomial expected value. To increase the robustness of the inference of changes in transcription and better help to anchor data from different samples for normalisation, the slope parameter for the main covariate (β_1_) was subject to a regularised horseshoe^89^ prior. The role of this prior is to impose a sparsity assumption on the gene-wise transcriptional changes; that is, most genes are not differentially transcribed. The overall distribution of the gene intercepts follows a gamma probability function. The statistical model is defined by the following joint probability density.

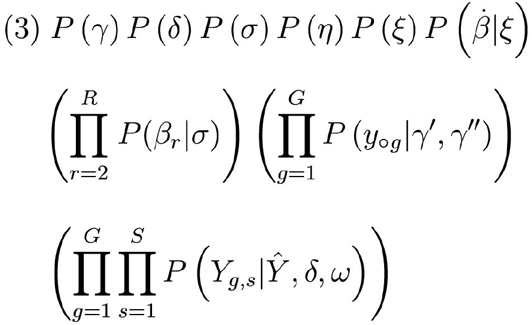

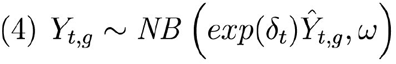

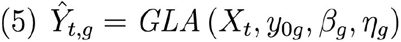

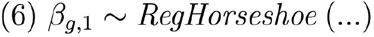

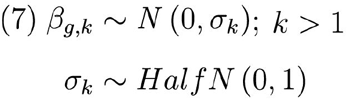

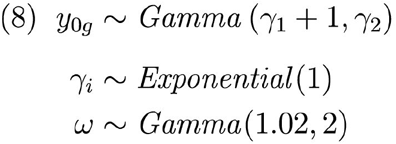

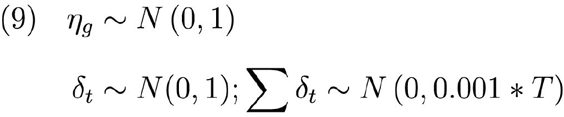

Where Y represents raw transcript abundance, *Ŷ* represents the expected values of transcript abundance and X represents the design matrix (with no intercept term and with scaled covariates). The regression function also includes β which represents the gene-wise matrix of factors (i.e., slopes excluding the intercept term), 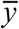 and η which represent the gene-wise y-intercept and the inflection point of the generalised reparameterised sigmoid function (Eq. 2), while γ represent the hyperparameters of 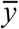. Other parameters of the negative binomial function are δ, which represents the normalisation factors; and ω, which represents overdispersion. The regularising prior (for imposing the sparsity assumption) over the covariate of interest β_1_ (first column of β) is defined by the hyperparameter list ξ^89^ (i.e., nu_local = 1; nu_global = 1; par_ratio = 0.8; slab_df = 4; slab_scale = 0.5), while σ represents the standard deviations of the other factors (in our case only the batch). The algorithm multidimensional scaling (MDS)^90^ was used to map the data in two dimensional space.

### Gene annotation

Each gene (g) was considered well fitted by the model if it had read counts outside the 95th percentile of the generated quantities for three or fewer samples (according to posterior predictive checks standards^20^). Among the well fitted genes, those for which the 0.95 credible interval of the posterior distribution of the factor of interest β_1g_ did not include the value 0 were labelled as differentially transcribed. The credible interval is a numerical range within which an unobserved parameter value falls within a certain probability. As distinct from common practices for frequentist models operating on confidence intervals and p-values, for this study the credible interval probability threshold was not altered for multiple hypothesis testing, consistently with common practices in Bayesian statistics^21^.

In order to interpret the inflection points over the CAPRA-S risk score (i.e., the point of maximum slope; at what stage of the disease a transcriptional change happens) covariate in a biologically meaningful way, the inflection point was adjusted to the log-scale. Considering that the lower plateau of our generalised sigmoid function was set to 0 (in order to limit the number of parameters needed to model it), the inflection point of the logarithm-transformed function is not defined. Therefore, we calculated the inflection point (*Ẋ*) of the log sigmoid forcing a plateau at 1 (i.e., log(0) = 1; Eq. 10; Fig. S7). This new inflection point can now be calculated as the value of the x-axis at half distance between zero and the upper plateau of the generalised reparameterised sigmoid function (Eq. 10).

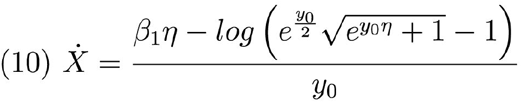

Genes were functionally annotated with gene ontology categories^22^ using BiomaRt^91^. Furthermore, genes were functionally annotated with the protein atlas database^92^ for identifying those that interface with the extracellular environment, encoding for cell-surface and secreted proteins. For a more in-depth analysis of possible interactions between cell types, we compiled a cell-type specific annotation database for cell-surface and secreted protein coding genes (Supplementary file 3).

### Differential tissue composition analyses

The differential tissue composition analysis is composed by: (i) a first deconvolution step that infers tissue composition from whole tissue gene transcript abundances based on reference transcriptional profiles of single cell types; and (ii) integrated beta regression on the inferred proportions. The use of a Bayesian inference framework (GitHub: stemangiola/ARMET@v0.7.1) allowed to transfer the uncertainty from (i) to (ii). The probabilistic framework Stan^88^ was used to encode the joint probability function of the model. The 0.95 credible interval of the posterior distributions was used for result interpretation. This algorithm bases the inference on 18 hierarchical cell-types defined by a collection of 250 curated publicly available transcriptional profiles (including BLUEPRINT^64^, ENCODE^65^, GSE89442^66^ and GSE107011^67^).

### Analysis of tumour microenvironment using multiplex immunohistochemistry

Slides (3μm) from formalin fixed and paraffin embedded (FFPE) tissue were taken from a total of 63 core biopsies of localised prostate cancer across 17 patients. A pathological evaluation was done to define tumour and surrounding benign tissue areas for each biopsy. The methodology for performing multiplex immunohistochemistry, cell type classification and localisation has been detailed by Keam et al. ^93^. Briefly, slides were deparaffinised and rehydrated with xylene and ethanol. The fluorochrome-coupled antibodies against human CD68 (macrophages and dendritic cells; DC), high molecular weight cytokeratin (HMWCK; epithelial basal cells), CD3 (T cells), CD20 (B cells), CD11c (dendritic cells), and PDL1. The dye DAPI was used for nuclei staining. Vectra 3.0 Automated Quantitative Pathology Imaging System, 200 slide (Perkin Elmer, MA) was used for imaging as detailed by Keam et al.^93^. The software HALO was used for cell segmentation and phenotyping. Stromal cells were defined with negative selection of all antibodies (DAPI positive), and with filtering by large size (cell area > 70) and highly elongated shape (ratio of largest dimension and smallest dimension > 2; 0.9 percentile; 0.9 percentile; Fig. S6).

Cell type proximity was quantified as the amount of cells within a radius of 20 cells sizes, from a selected cell, averaged per tissue area (5 cell size units) for smoothing and avoiding information duplication due to tight cell clusters. Cell relative size was calculated at 15 units, as the observed median length units in the data coordinate system. The statistics were summarised at the biopsy level. When the distance between two cell types was measured, only the biopsies including both cell types were selected. The robust regression analyses were performed using the R heavy package^94^ on log transformed proximity measure. The co-proximity analysis between epithelial basal cells and PDL1+ macrophages and T cells, was performed at single cell level (averaged by tissue area of 5 cell size units) calculating the proximity on a radius of 50 relative cell sizes for ensuring a good coverage of both T cells and PDL1+ macrophages, and decrease sparsity. Only the epithelial basal cells in immune rich areas (with > 5 neighbour T cells) were considered.

## Notes

### Competing Interest Statement

The authors have declared no competing interest.

