## Supplementary Material for "Dissection of prostate tumour, stroma and immune transcriptional components reveals a key contribution of the microenvironment for disease progression"

### Supplementary results

#### Quality control

After trimming and filtering, the sequenced reads of all samples achieved a Phred quality score above 28. The sequencing output ranged from 70 million (for sample 2C; Fig. S2) to 1 million reads (for sample 4C; Fig. S2). Most samples (n = 41) had more than 80% of reads uniquely mapped to the hg38 reference genome, with an exonic rate in the range of 50 to 70%. Overall, myeloid samples were characterised by a lower sequencing and mapping coverage compared to the other three cell types.; For all samples, a positive association was observed between CAPRA-S risk score and (i) number of mapped reads, as well as (ii) number of mapped reads to exons (Fig. S8). This association was partly due to an abundant amount of residual bacterial and viral genomic content especially for the surrounding benign and low-grade cancers (Fig. S8).

### Supplementary tables

| Patient | ID Age | Serum PSA (ng/ml) | Gleason Grade | T-stage | CAPRA |
|---|---|---|---|---|---|
| RA014 | 57 | 11 | 3 | 1c | 3 |
| RA020 | 62 | 7.9 | 3 | 2b | 2 |
| RA025 | 57 | 80 | 3 | 3b | 5 |
| RA030 | 70 | 8.5 | 3 | 2b | 2 |
| RA031 | 66 | 6.4 | 5 | 2a | 5 |
| RB010 | 64 | 20 | 3 | 2b | 3 |
| RB011 | 68 | 10.9 | 2 | 2a | 3 |
| RB023 | 64 | 3.7 | 3 | 1c | 1 |
| RB032 | 76 | 27.7 | 4 | 1c | 6 |
| RB037 | 73 | 7.2 | 5 | 2b | 5 |
| RB040 | 67 | 10.3 | 5 | 3b | 7 |
| RB050 | 66 | 7.3 | 2 | 1c | 2 |
| RB051 | 65 | 4.1 | 2 | 1c | 1 |

| | | | | | |
|---|---|---|---|---|---|
| RB052 | 70 | 6.1 | 4 | 2a | 4 |
| RB053 | 69 | 10.1 | 2 | 2b | 3 |
| RB056 | 66 | 4 | 3 | 1c | 1 |

**Table SX.** Summary of the clinical characteristics of the subjects included in the multiplex immunohistochemistry analysis of primary prostate tumour biopsies.

### Supplementary figures

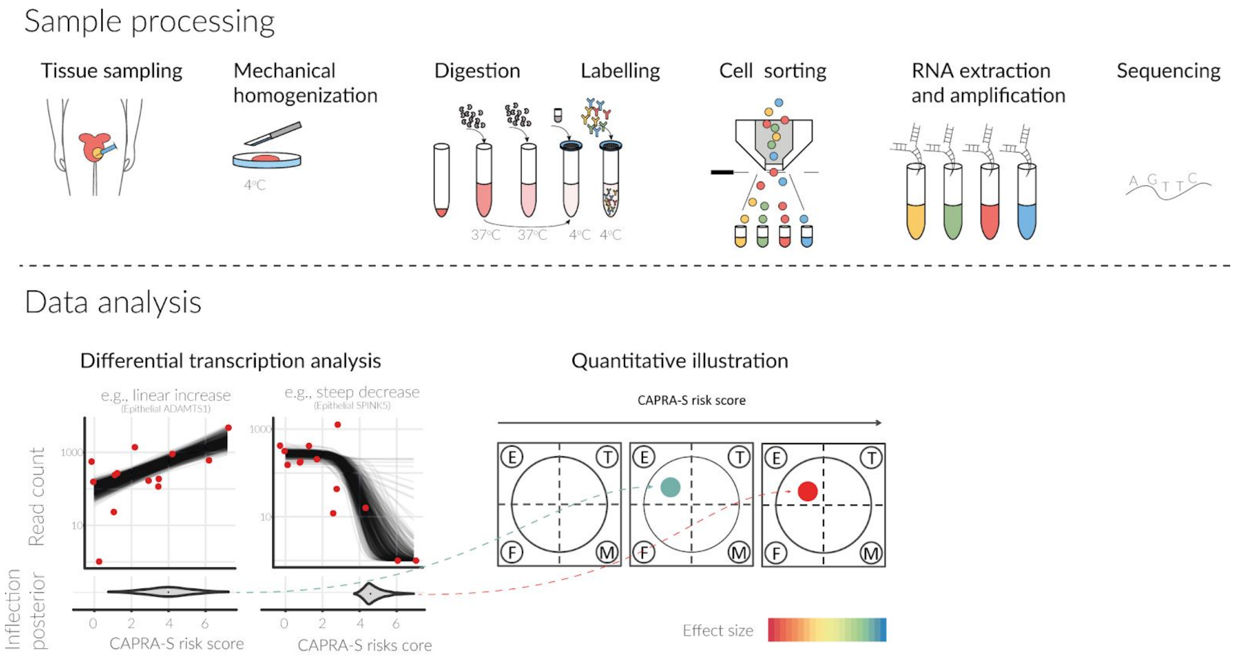


**Figure S1.** Diagram of the experimental and computational analysis pipeline.

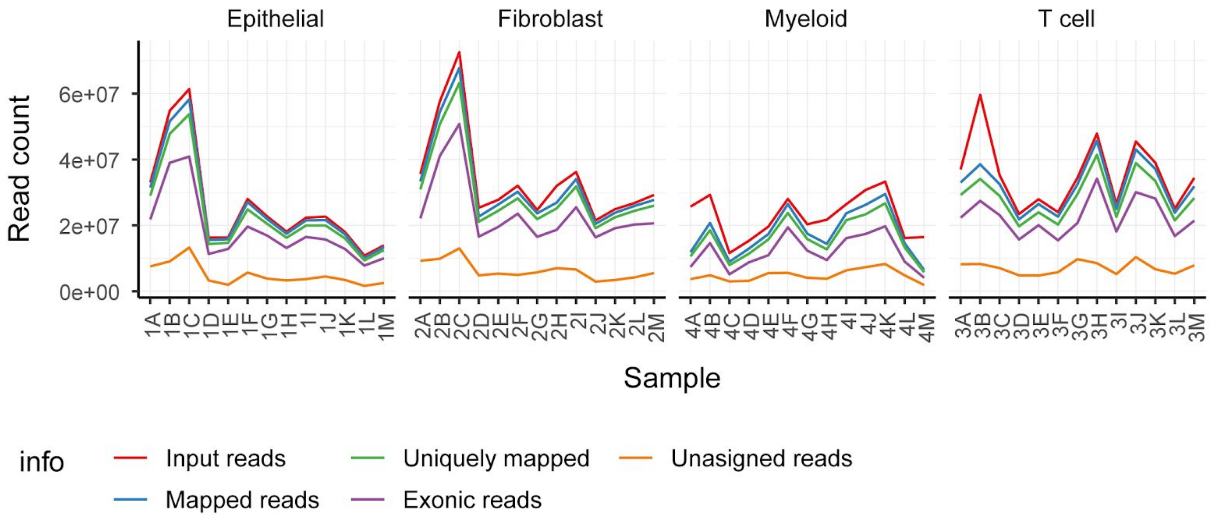


**Figure S2.** Mapping statistics for each sample, grouped by cell-type.

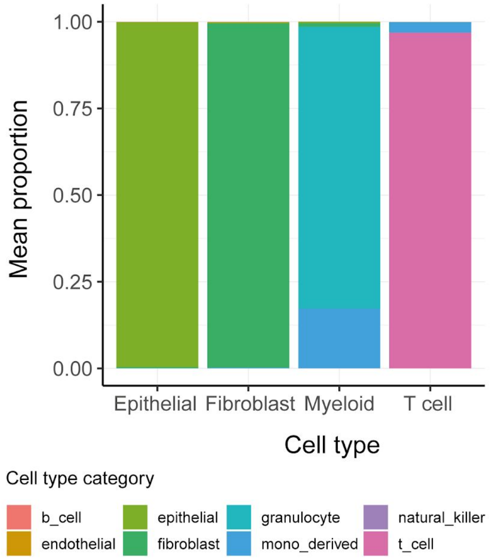


**Figure S3.** Bar-plot of cell-type composition for the four enriched cell types (E = epithelial; F= fibroblast; M = myeloid; T= T cell), Inferred by Bayesian inference deconvolution model.

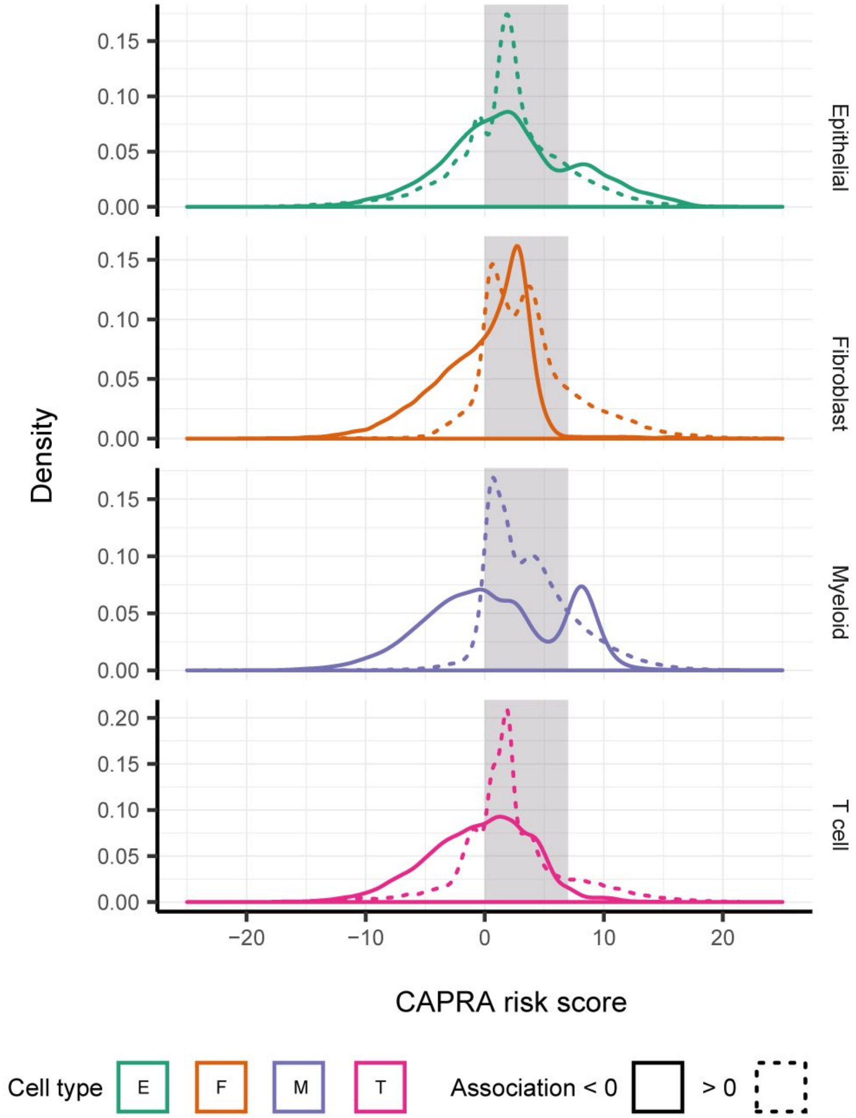


**Figure S4.** Distribution of the adjusted inflection points along the CAPRA-S risk score, across cell types. The grey shade represents the range of CAPRA-S risk score in the 13-patient cohort. Inflection points outside that range represent exponential-like trends (either increasing or decreasing) that did not approach a plateau within the CAPRA-S risk score range.

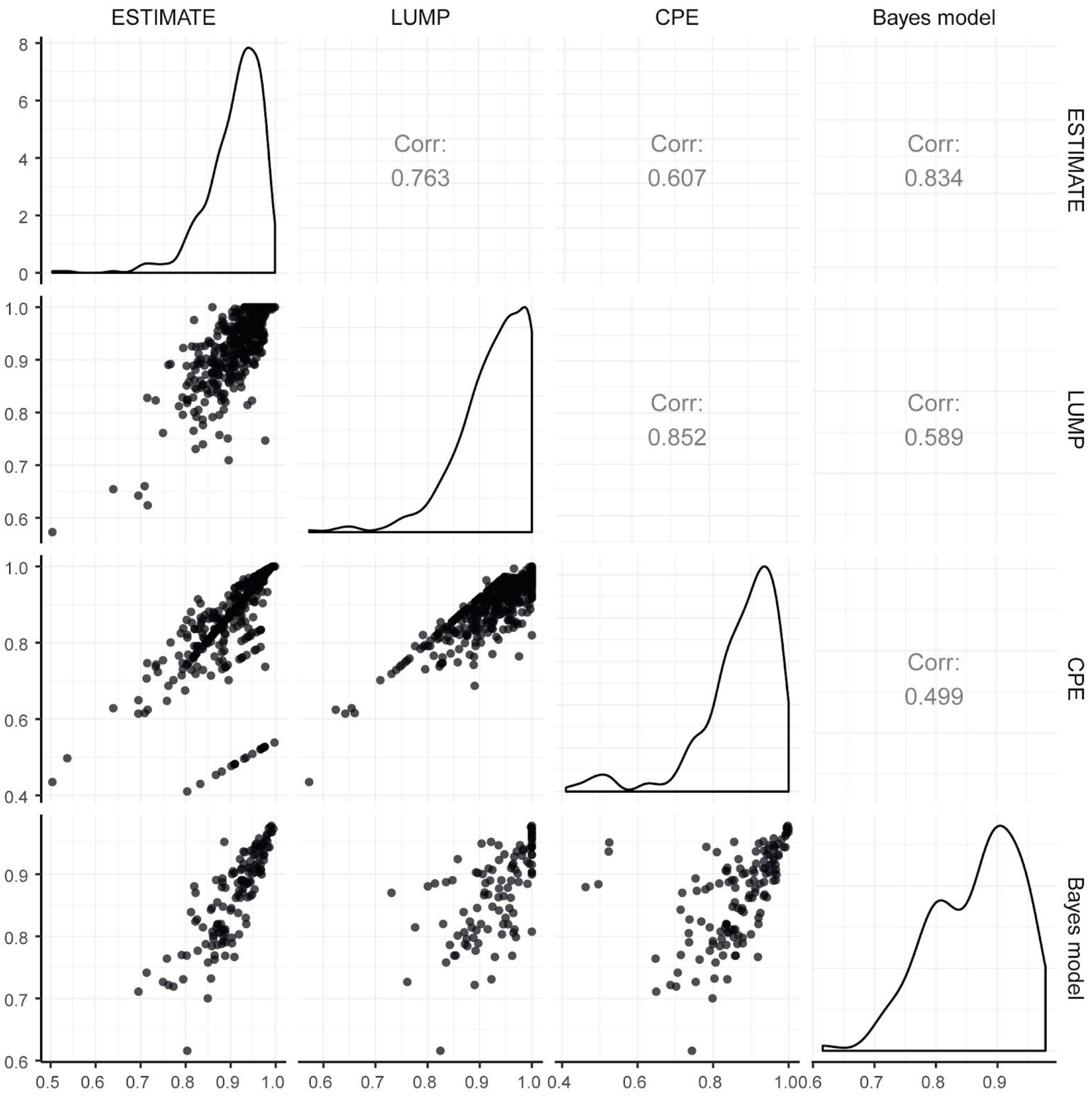


**Figure S5.** Pair plot showing the comparison between the inference of epithelial component in TCGA samples for our Bayesian inference model and gold standard tumour purity analysis[68].

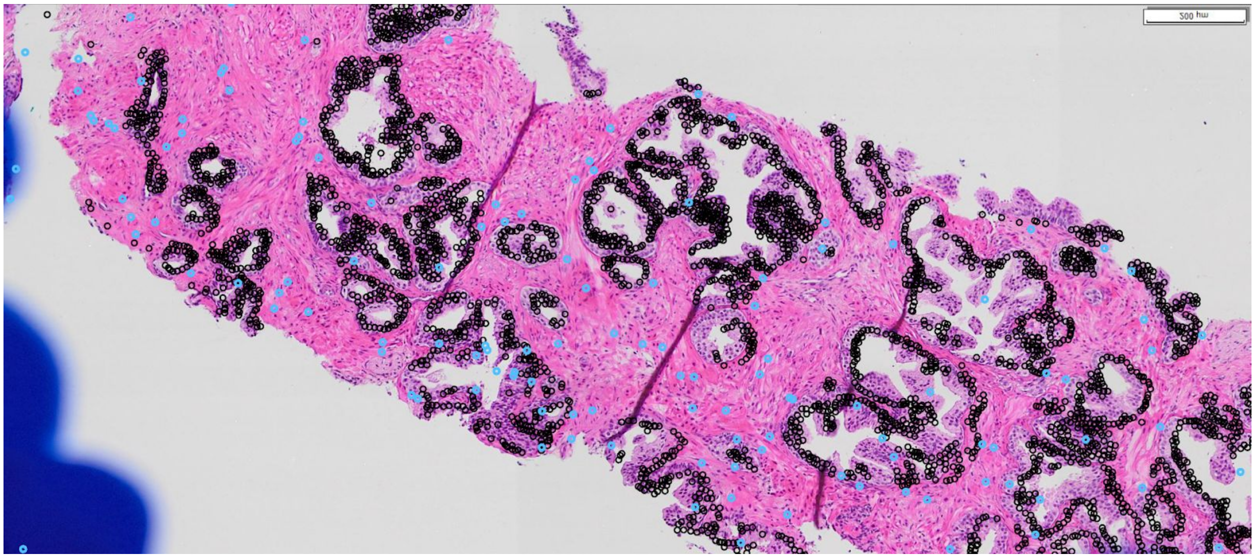

**Fig S6.** Example of stromal cells selection (blue circles). The black circles represent the HMWCK positive cells (epithelial basal). Stromal cells were labelled being negative for all markers, being DAPI positive, being larger than units 70 and being a highly elongated shape, with a ratio of largest dimension and smallest dimension > 2. These simple and stringent criteria lead to a highly specific but lowly sensitive selection. Specificity was prioritised over sensitivity considering the high number of stromal cells.

A

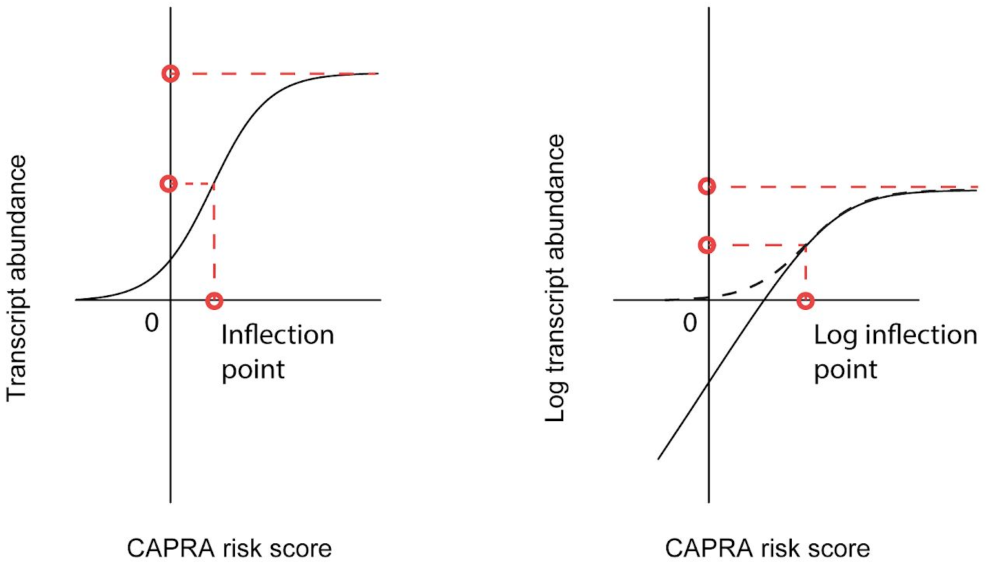


B

$$\text{Inflection of } log\left(GLA\left(\ldots\right)\right) = \frac{log\left(\text{ upper plateau of } GLA\left(\ldots\right)\right)}{2}$$

$$= log\left(y^{\circ}\right) + log\left(1 + e^{\eta\dot{\beta}}\right) - log\left(1 + e^{\eta\dot{\beta} - \dot{X}\dot{\beta}}\right) = \frac{log\left(y^{\circ}\right) + log\left(1 + e^{\eta\dot{\beta}}\right)}{2}$$

$$\dot{X} = \frac{\dot{\beta}\eta - log\left(e^{\frac{y^{\circ}}{2}}\sqrt{e^{y^{\circ}\eta} + 1} - 1\right)}{y^{\circ}}$$

**Figure S7. A** — Illustration of the strategy for the identification of the inflection point of the sigmoid function (left-panel) in log-scale (right-panel). The challenge is that in order to limit the number of parameters that the model needs, the lower plateau of the generalised sigmoid function was set to zero. Therefore, the inflection point of the logarithm transformed function is not defined (i.e., negative infinite). In order to define it, the lower plateau of the logarithm transformed function was also set to 0 (rather than -Inf). **B** — Formulae used to perform the transformation.

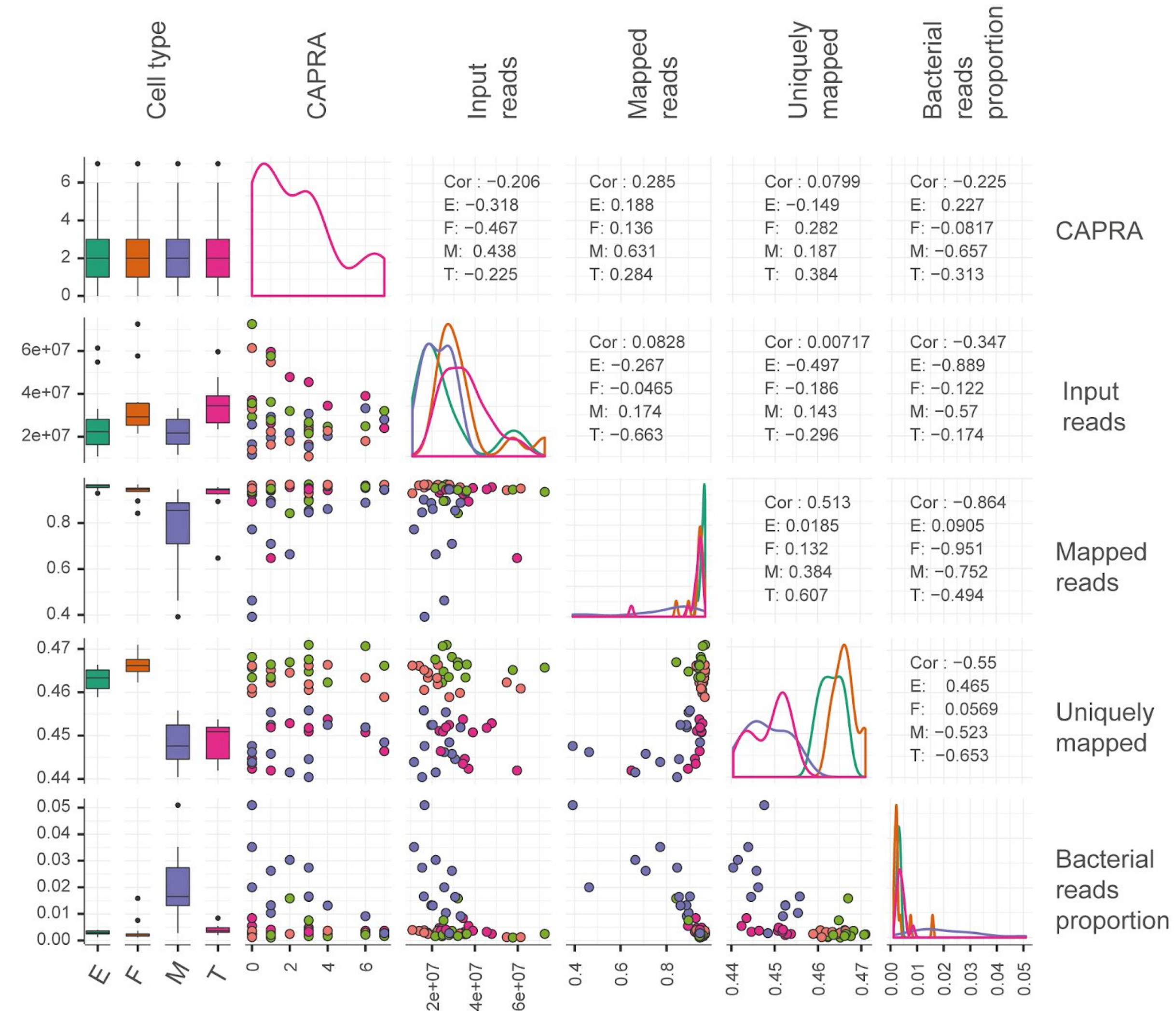


**Figure S8.** Pair plot showing the relations among sequencing/mapping statistics and CAPRA-S risk score, stratified by cell type (E = epithelial; F = fibroblast; M = myeloid; and T = T cell).
